# The effect of insect cyanoglucosides on predation by domestic chicks

**DOI:** 10.1101/662288

**Authors:** Márcio Zikán Cardoso

## Abstract

Cyanogenic insects release cyanide and other byproducts that are thought to make them unpalatable to would be predators. In fact, aposematic *Heliconius* butterflies and *Zygaena* moths are rejected by vertebrate predators. Nevertheless, there have been few studies testing the effect of cyanoglucosides on vertebrate predation. Here I report tests undertook with captive domestic chicks in order to evaluate the effect of two chemically diverse cyanoglucosides, linamarin and prunasin. In insects as well as plants, cyanoglucosides are stored in vacuoles and are enzymatically broken down when the tissue is disrupted as in the case of a predator attack. Linamarin is an aliphatic that releases cyanide and acetone upon breakdown, while prunasin is an aromatic cyanoglucoside that releases cyanide and benzaldehyde. Using concentrations that mimicked the average concentration of a *Heliconius* butterfly, supplemented by test with higher concentrations, I ran a series of trials with naïve chicks by offering prey laced with chemicals. I categorized prey acceptance and compared the behavior of the predators towards test and control larvae. Prey coated with cyanide and benzaldehyde were significantly rejected by the birds, while acetone did not elicit avoidance behavior. Intact cyanoglucosides apparently were not detected by the predators, presumably because of fast ingestion time or lack of enzymes to breakdown cyanoglucosides. The rejection of cyanide laced prey confirm the protective nature of cyanoglucosides against a vertebrate predator. Additionally, the rejection of the pungent but not toxic benzaldehyde suggests that some species that store aromatic cyanoglucosides could be detected via smell as well by taste. These results provide support for cyanoglucosides as defensive chemicals of aposematic lepidopterans and related arthropods.

## Introduction

Cyanogenic glycosides (cyanoglucosides) are found in arthropods such as millipedes, centipedes and insects (notably moths and butterflies) and are considered defensive chemicals against predators (Zagrobenly et al 2008, 2018). Cyanogenic glycosides release hydrogen cyanide (HCN) when tissues containing them are ruptured. In human societies, this phenomenon accounts for acute and chronic cyanide toxicity and is a source of health concerns for populations that rely on cyanogenic cassava plants for consumption (Jørgensen et al 2005). Since action of cyanide in biological systems is well documented, it is reasonable to expect that cyanogenic glycosides are toxic to predators as well.

Two of the most studied cyanogenic lepidopterans are the aposematic moth family Zygaenidae (burnet moths) (Zagrobelny et al 2008) and Heliconiinae butterflies (Nahrstedt 1988, Castro et al 2019), many of which have been shown to be unpalatable to birds and other predators (Brower et al 1963, Boyden 1976, Chai 1986, Pinheiro & Campos, 2019). The insectivorous jacamar (*Galbula ruficauda*) discriminates between palatable (*Dryas iulia* and *Agraulis vanillae*) and unpalatable Heliconiinae (*Heliconius*) butterflies (Chai 1990), which is consistent with differences in cyanoglucoside accumulation patterns in these species (Nahrstedt & Davis 1983, Cardoso & Gilbert 2013).

The variety of chemical structures of the cyanoglucosides found in arthropods is lower than that found in plants (Zagrobelny et al 2018). The aliphatics, linamarin and lotaustralin (Figure 1), are commonly found in lepidopterans, usually with the former being more abundant. Additionally, some Heliconiines may also contain the more complex cyclopentenoid cyanoglucosides (Castro et al 2019). Aromatic cyanoglucosides (Figure 1) are more commonly found in millipides and relatives, although a few insects also produce them (Zagrobelny et al 2018).

**Figure 1.**
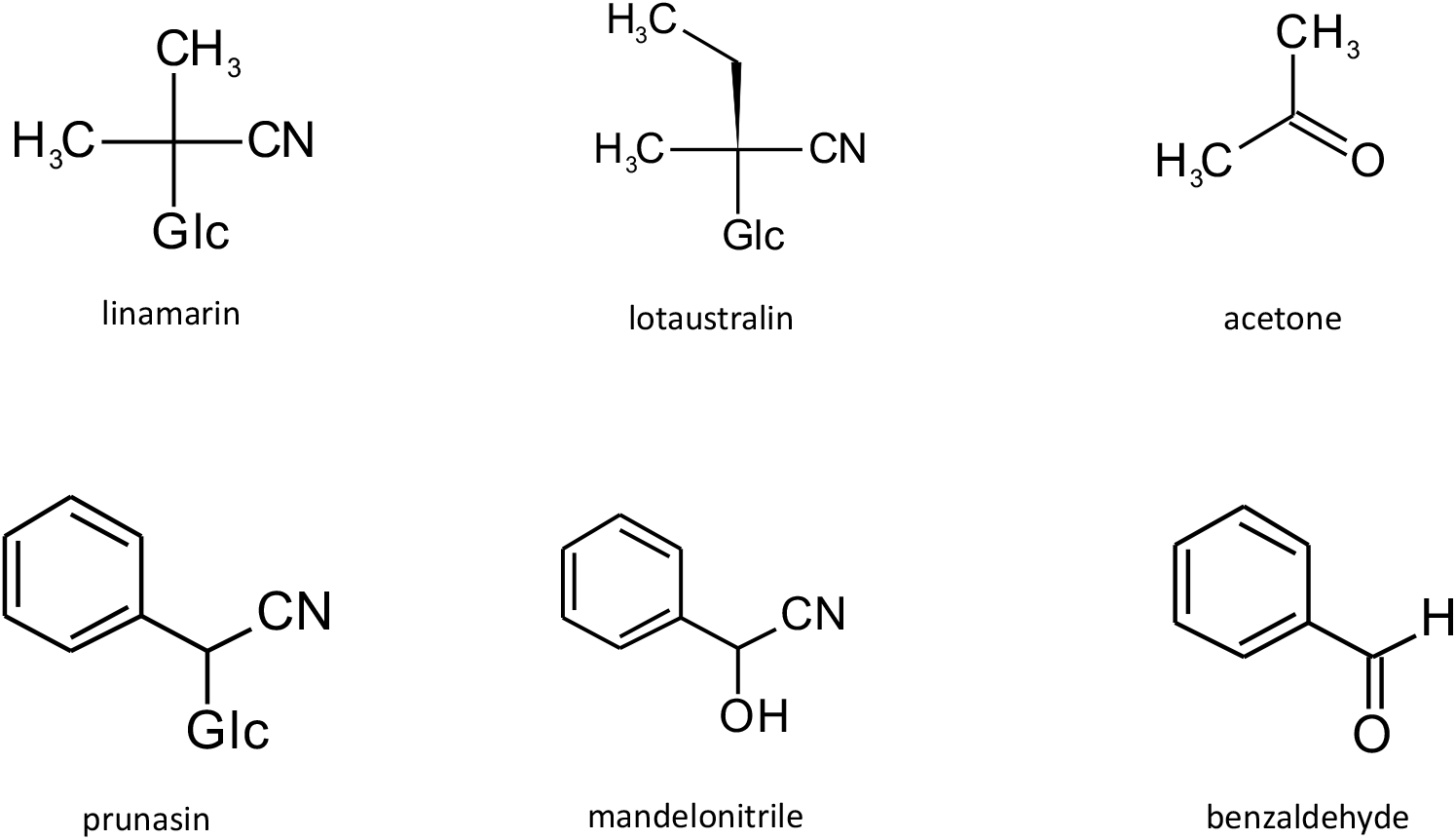
Chemical structure of selected cyanoglucosides. Linamarin and lotaustralin are aliphatics commonly found in lepidopterans, usually with the former being more abundant. Acetone is one of the breakdown products of linamarin. The aromatic cyanogenic glycoside, prunasin, is a main component in many cyanogenic plants and is chemically related to mandelonitrile, which is present in many arthropods. Benzaldehyde is released upon breakdown of prunasin.

The enzymatic breakdown of cyanoglucosides releases not only HCN but also other byproducts that are structure-specific, such as the case of acetone from the aliphatic linamarin, and benzaldehyde from the aromatic cyanoglucosides prunasin and mandelonitrile (Figure 1). Thus, the defensive potential of cyanoglucosides may reside not only on the release of HCN but also on some of these byproducts.

Although *Heliconius* unpalatability is a classic theme in evolutionary ecology, there have been few studies that tested whether the cyanogenic compounds they store can make them unpalatable. In fact, a literature search has found only one study that directly tested the effect of a cyanoglucoside on a vertebrate predator (Muhtasib & Evans 1988). This study found that linamarin, the major cyanoglucoside in lepidopterans, elicit avoidance in quails at high concentrations. However, when applied together with histamine, which is also found in *Zygaena*, the effects were much more pronounced, suggesting that a synergistic interaction could exist. Using ants as test subjects, Peterson (1986) tested which of the breakdown products from prunasin (present in the gut contents of a *Prunus* herbivore, *Malacosoma*) elicited rejection in them. Curiously, the toxic cyanide moiety was not detected by the predators but another breakdown product, benzaldehyde, although not toxic, was rejected by them. These results suggest not only that cyanoglucosides are deterrent to predators, but also that olfactory cues provided by breakdown products could be used by predators to detect cyanide in the prey. Given the general lack of tests on the palatability of cyanoglucosides to vertebrate predators, I set out to test the response of captive chicks to two cyanoglucosides and their breakdown products. I tested two cyanoglucosides that differ in structure by using linamarin (an aliphatic) and prunasin (an aromatic). Prunasin is not common in animals, but it is an aromatic cyanoglucoside chemically related to mandelonitrile, which is found in many arthropods and also releases benzaldehyde upon breakdown.

## Material and methods

### Cyanoglucosides

I evaluated the palatability of two cyanoglucosides, linamarin and prunasin, and of their byproducts (Figure 1) to captive naïve domestic chicks. Both substances release hydrogen cyanide upon degradation and I used NaCN as a substitute for the HCN gas in the experiment. The two other breakdown products tested were the volatiles acetone (from linamarin) and benzaldehyde (from prunasin). All chemicals were obtained from commercial suppliers, except linamarin which was obtained from root extracts of a highly toxic local variety of *Manihot esculenta* (Euphorbiaceae). TLC analysis of the extract showed that lotaustralin was also present in some samples. Lotaustralin usually co-ccurs in all insects that store aliphatic cyanoglucosides.

### Palatability assays

Trials were conducted using 7-day old naïve chicks as predators. Chicks were obtained from a local hatchery in groups of 10-20 individuals and kept together in an enclosure for 2-3 days (80 × 80 × 90 cm). They were then separated and kept in screened individual cages (28 × 25 × 26 cm) where palatability trials were performed. Chicks were provided water *ad libitum* and fed a commercial brand of chick feed twice a day. Food was removed from the cage one hour before the trial began. All animals were kept in a climate-controlled room (temperature: 25 ± 3 °C) with natural as well as artificial light. To familiarize the chicks with the experimental protocol I offered 3-6 larvae of the experimental prey, *Hermetia illucens* (Diptera), for three days prior to the trials.

Palatability assays consisted on presenting a 3-larvae sequence to the birds. The series started with a control larva. If accepted, then an experimental prey was offered, followed by a second control larva. If a bird rejected the first control offered, it was not included in the experiment (this only occurred in six birds out of a total of 129 animals we tested). Therefore, first controls were, by definition, always eaten. We used an average of 14 (± 5 sd; min = 8, max = 23) birds per experiment. Larvae were individually offered in a Petri dish and left in the cage for up to three minutes or less, if the predator ingested it. Bird responses were categorized as approach, attack, ingestion and were timed during trials. Test larvae were coated with 10 μl of the test cyanoglucoside or byproduct and left to dry prior to being offered to the bird. Control prey were coated with solvent only. In the case of the volatiles, we offered prey immediately after application of the chemicals and controls were larvae devoid of the volatile.

Bird response was categorized either as acceptance (predator attacks, handles and ingests prey) or rejection (predator does not attack, or predator attacks and releases prey, no ingestion occurs). I used contingency tables to test for an association between prey type (control vs. experimental) and bird response (accept vs. reject) and analysis of variance to analyze time to ingestion. All analyses were performed in JMP 5.0.1 (SAS Institute 2002).

Because the goals of this study were to analyze *Heliconius* defenses, I chose to work with the cyanoglucoside concentration normally found in this group. Based on Nahrstedt & Davis (1983), the average cyanide concentration in *Heliconius* is near 2 μmol/individual or 58 μmoles/gram dry weight. Thus, this average concentration was used in the trials, adjusting to the lower larvae dry weight (56.4 mg ± 10.31, N = 20). In some trials where enough chemicals were available, I also tested larvae with twice the average concentration.

## Results

### Overview of the experiments

I tested 123 birds in nine palatability trials. Overall, time to prey ingestion was not different across trials (F_8,330_ = 1.28, P = 0.25), indicating that birds were consistent among themselves, even when time to ingestion was affected by prey type (see below). The average time to ingestion was very fast, 10.6 ± 18.1 seconds (min = 1, max = 180 seconds, N = 339). Ingestion occurring in 10 seconds or less accounted for 75% of the observations.

### Cyanogenic glycosides

I ran 4 trials with the cyanoglucosides, three with linamarin and one with prunasin. With linamarin, the average concentration was used in 2 trials, and higher concentration was used in the third trial. Regardless of type and concentration, there was no noticeable negative effect of the intact cyanoglucosides on the birds. In the trials using average linamarin concentration, for instance, only 1 experimental prey was rejected, a rejection rate of 8.3%. A similar response was observed in the high concentration trial (rejection rate = 6.7%) (Figure 2). In the trials with prunasin, no experimental larvae were refused (N = 15 experimental larvae). In all cases bird behavior was not significantly different from control prey (Figure 2). Moreover, time to prey ingestion, combining the four trials, was not affected by prey type (F_2,157_ = 0.64, P = 0.53).

**Figure 2.**
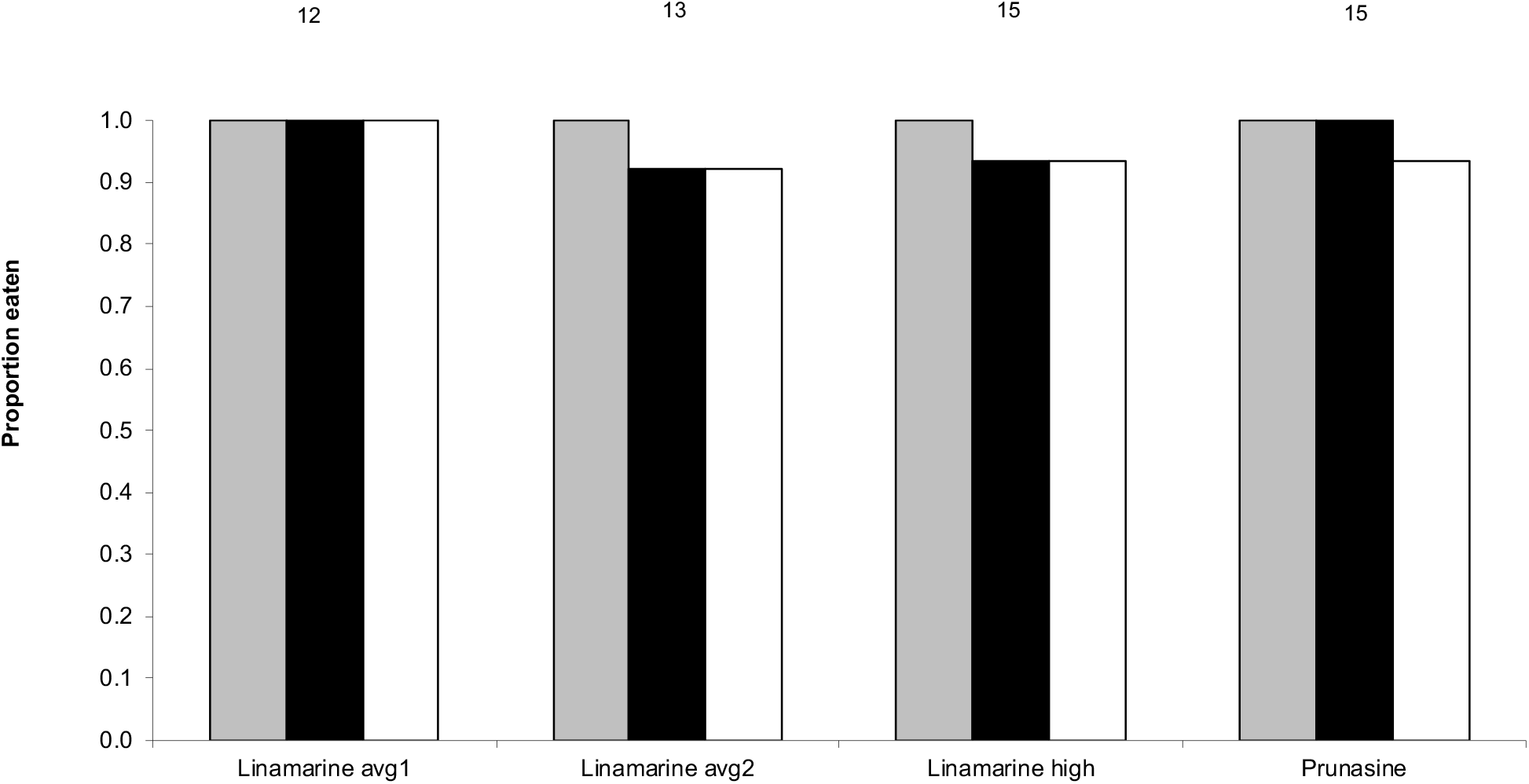
Proportion of prey ingested in the trials using intact cyanoglucosides. Birds were presented with three larvae in sequence, control 1 (grey bars), experimental (black bars) and control 2 (white bars). Concentrations mimicked the concentration of adult *Heliconius* butterflies (labeled as *avg* in graph) or were two times higher (*high* label). A number after a chemical name refers to a replicated experiment, while numbers above bars depict the number of larvae tested per treatment per experiment.

### Breakdown products: Volatiles ‒acetone and benzaldehyde

Acetone is the breakdown ketone from linamarin. To the experimental birds, presence of the acetone, even when larvae were literally drenched in it, did not affect acceptance (Figure 3). In the nine trials performed, no larva was rejected. Although eventually all prey were ingested, there was a tendency for birds to take longer to ingest acetone coated prey than control prey (F_2,24_ = 2.739, P = 0.085).

**Figure 3.**
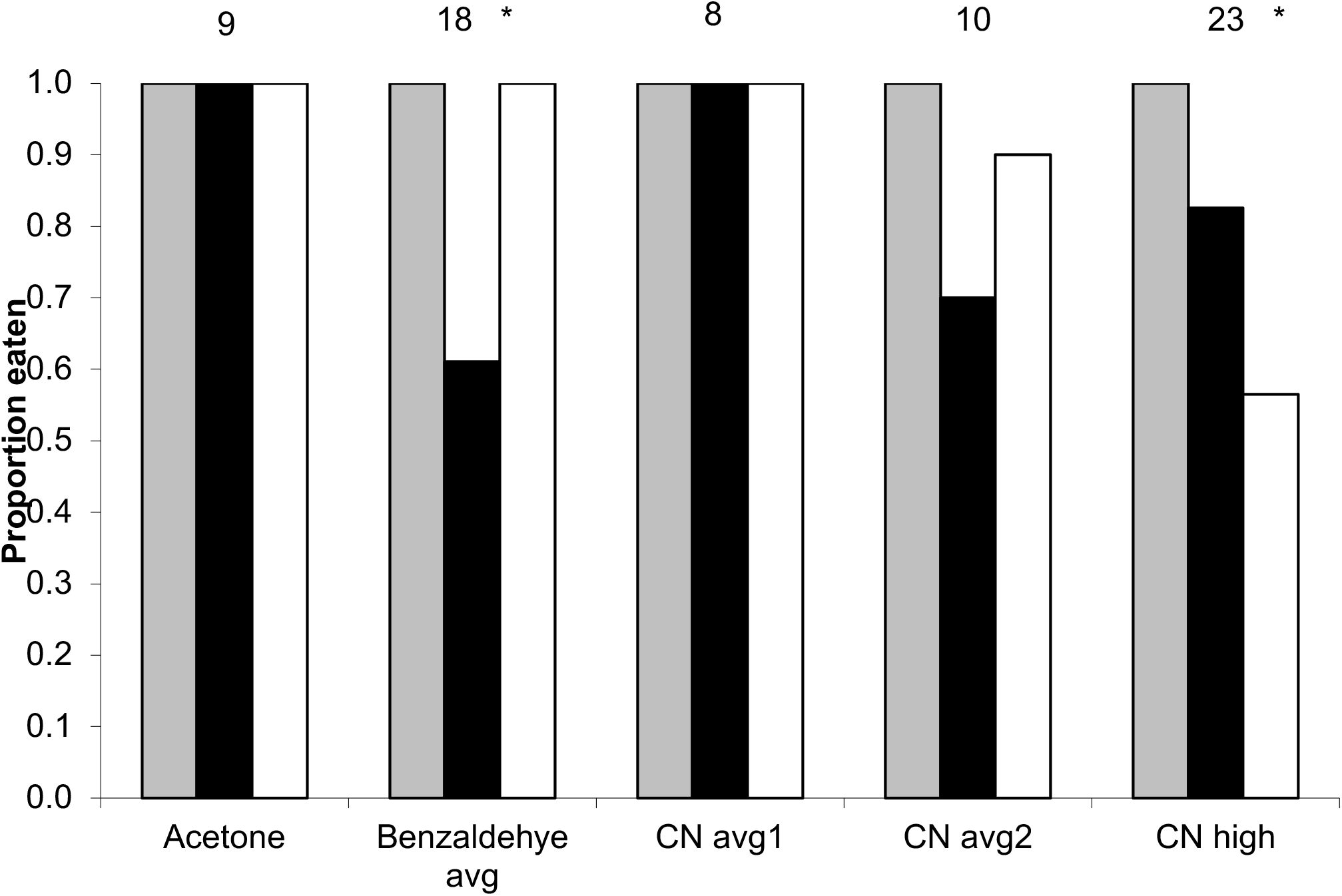
Proportion of prey ingested in the trials using breakdown products. Sodium cyanide (NaCN) was used to simulate HCN. Birds were presented with three larvae in sequence, control 1 (grey bars), experimental (black bars) and control2 (white bars). Concentrations mimicked the concentration of adult *Heliconius* butterflies (labeled as *avg* in graph) or were two time higher (*high* label). Numbers after name refer to a replicated experiment. Numbers above bars represent the number of larvae tested per treatment per experiment. *= P < 0.05, CN = cyanide (NaCN).

Application of benzaldehyde, the volatile released by degradation of prunasin, significantly affected bird behavior (G = 17.59, P = 0.0002) (Figure 3). There was an increase in the rejection rate of experimental prey, nearing 40%. Moreover, time to ingestion was also affected (F_2,44_ = 6.91, P = 0.002), with benzaldehyde coated prey taking longer to be ingested (average = 23.3 seconds) than control prey (average = 3.9 seconds). Additionally, the ingestion of the second control tended to occur later than ingestion of first control, suggesting a carry-over effect of the experimental prey on predator behavior.

### Cyanide

Application of sodium cyanide elicited rejection behavior in the chicks, most notably in the high concentration trial (Figure 3). I conducted three trials, two using the average butterfly concentration and one using the higher concentration. No larvae were rejected in the first average concentration trial (8 predators, 24 larvae). In the second trial, however, 30% of the experimental prey were not eaten (N = 10 larvae), which led to a near significant association between larval type and behavior (G = 4.84, P = 0.089). No effect of time to ingestion was found in these two trials (F_2,47_ = 0.858, P = 0.431). When the high concentration was used in the experimental prey, birds began rejecting not only the experimental but also the second control prey (Figure 3). In fact, the combined rate of rejection was 30.4% (10 and 4, control 2 and experimental, respectively), a significant effect of bird behavioral change (G = 16.88, P = 0.0002). Time to ingestion was not significantly affected in this experiment (F_2,52_ = 1.998, P = 0.146).

## Discussion

The storage of cyanoglucosides in arthropods is thought to serve a protective function against predators (Zagrobelny et al 2008, 2018), yet few studies have tested this hypothesis on vertebrate predators, the main natural enemies of aposematically colored lepidopterans (Pinheiro & Cintra 2017). Here, I show that defense based on cyanoglucoside is effective against such a predator.

Application of sodium cyanide elicited negative responses from the birds. This effect was seen at higher concentrations, showing that predators are capable of detecting cyanide and rejecting prey laced with it. In addition, a breakdown product of aromatic cyanoglucosides, the volatile benzaldehyde, also elicited aversive behavior in the experimental predators. Benzaldehyde gives off a clear smell and is the main component of bitter almond oil. In fact, experiments with almond oil show that it can potentialize predator learning (Roper & Marples 1997). Similarly, Peterson (1986) found that ants avoided prey coated with benzaldehyde. Consequently, aromatic cyanoglucosides can be detected by predators by both of their breakdown products. Contrarily to the results with benzaldehyde, the volatile acetone from the aliphatic cyanoglucoside linamarin, did not seem to affect predator behavior. Larvae drenched with the substance were ingested with no obvious repulse by the predator.

It was also clear that the predators were not affected by intact cyanoglucosides. In real prey, immediate enzymatic action would release breakdown products. For this to occur in this experiment, cyanoglucosides would have to be degraded by the predator’s own set of glucosidases which are probably not specific for the experimental cyanoglucosides (Pentzold et al 2017). Even if this occurred, I suspect that the rapid consumption time (< 10 seconds on average) and the small prey size did not allow enough time for a gut reaction to be detected. Since prey handling may allow the predator to assess texture as well as taste, this finer assessment of prey was lacking in the “predators”.

Prior to this study, there were few direct tests of cyanoglucoside palatability in vertebrates. Muhtasib & Evans (1987), found aversive response in quails offered water with linamarin concentrations that were equal to or higher than the concentration in adult moths, not noticeably higher than that of *Heliconius* butterflies (Davis & Nahrstedt 1981, Nahrstedt & Davis 1983), which suggests that the intact cyanoglucosides can be detected by a predator. It would be instructive to know the detection thresholds of regular butterfly predators which are probably more fine-tuned, given its regular exposure to cyanogenic prey (Pinheiro & Campos 2019). One important aspect from Muhtasib & Evans (1987) work is that a lower linamarin concentration was effective when combined with additional defensive compounds such as histamine. In the *Heliconius* system it is not known whether other chemicals could be used as subsidiary effectors (Zagrobelny et al 2018, Castro et al 2018).

Brower (1984) once proposed that harmful defensive chemicals that could be detected by predators (e.g. bitter taste of alkaloids) should be called type I defensive chemicals, while chemicals that were used as proxies for a more toxic compound but were of no effect themselves should be called type II. Could the defensive systems of cyanogenic organisms be based on a signal that combines smell (type II) and taste (type I)? The evidence suggests so. In contrast to the moderate response by the chicks, a specialized predator reacts in a non-hesitant fashion to *Heliconius* taste. For instance, even when free flying *Heliconius* is captured by a jacamar, they are soon released without significant harm (Langham 2004, Pinheiro & Campos 2019), which clearly shows that these insects are unpalatable.

Although we now have a better understanding of cyanoglucoside defenses, further work is still needed. For instance, it would be instructive to know if more complex cyanoglucosides such as those sequestered by specialized *Heliconius* (Castro el al 2019) provide better protection against predators, and to evaluate if the accumulation of cyanoglucosides by palatable heliconiine species such as *Dryas* and *Agraullis* suggest additional roles for these compounds (Cardoso & Gilbert 2013, Zagrobelny et al 2018). These will allow a better mechanistic understanding of the defensive chemicals that permitted the evolution of aposematism in *Heliconius* and *Zygaena*.

## Acknowledgments

I would like to dedicate this study to Professor José Roberto Trigo (1956-2017), head of the Chemical Ecology Lab at the University of Campinas. Dr Trigo provided space, logistical support, and the chemicals needed for this study, to which I am grateful. Sadly, Dr Trigo passed away before seeing the final draft of this work. I would also like to thank FAPESP for a postdoctoral fellowship for the author and laboratory grants to JRT. I thank M.A. Foglio (CPQBA, Unicamp) for providing linamarin samples and the Agronomical Institute in Campinas for the cassava material. Special thanks are due to J. Paiva for assistance with chemical analysis, P. R. Guimarães for helping with the palatability experiments, and Érika Castro with the drawing of chemical structures. I would also like to thank Mika Zagrobelny and Érika Castro for critically reviewing the manuscript.

